# Integration of A Nitrate-Related Signaling Pathway in Rhizobia-Induced Responses During Interactions with Non-Legume Host Arabidopsis thaliana

**DOI:** 10.1101/2020.09.08.287219

**Authors:** Sebastian T. Schenk, Elisabeth Lichtenberg, Jean Keller, Pierre-Marc Delaux, Thomas Ott

## Abstract

Nitrogen (N) is an essential macronutrient and a key cellular messenger. Plants have evolved refined molecular systems to sense the cellular nitrogen status. Exemplified by the root nodule symbiosis between legumes and symbiotic rhizobia, where external nitrate availability inhibits the interaction. However, nitrate also functions as a metabolic messenger, resulting in nitrate signaling cascades which intensively cross-talk with other physiological pathways. NIN (NODULE INCEPTION)-LIKE PROTEINS (NLPs) are key players in nitrate signaling and regulate nitrate-dependent transcription. Nevertheless, the coordinated interplay between nitrate signaling pathways and rhizobacteria-induced responses remains to be elucidated. In our study, we investigate rhizobia-induced changes in the root system architecture of the non-legume host Arabidopsis in dependence of different nitrate conditions. We demonstrate that rhizobia induce lateral root growth, and increase root hair length and density in a nitrate-dependent manner. These processes are regulated by AtNLP4 and AtNLP5 as well as nitrate transceptor NRT1.1, as the corresponding mutants fail to respond to rhizobia. On a cellular level, *NLP4* and *NLP5* control a rhizobia-induced decrease in cell elongation rates, while additional cell divisions occurred independent of *NLP4*. In summary, our data suggest that root morphological responses to rhizobia, dependent on a nutritional signaling pathway that is evolutionary related to regulatory circuits described in legumes.

## Introduction

Nitrogen, as a plant macronutrient and limiting factor for plant growth, is highly demanded in agriculture. Thus, organic and synthetic nitrogen fertilizers are essential for productive farming (Vitousek *et al*., 2009). The advancement of fertilizer usage and the need for sustainable solutions impulses research in two crucial perspectives (i) the molecular investigation of nitrate sensing/signaling of plants, to improve and understand nitrogen use -efficiency and -processing and (ii) rhizospheric bacteria, that are important for the mineralization of nitrogen in the soil, but could also associate with plants and influence plant nitrogen consumption and uptake.

In this context, nitrogen-fixing (diazotrophic) bacteria play a key role in making dinitrogen gas available for plants in the rhizosphere (Carvalho *et al*., 2014; Rosenblueth *et al*., 2018). Besides the endosymbiotic nitrogen fixation with rhizobia species and their specialized legume host, there are several free living, associated-, and/or endophytic- rhizobacteria that interact with non-legume plant hosts and promote a better plant growth (Hardoim *et al*., 2008; Van Deynze *et al*., 2018). These mutualistic bacterial interactions facilitate biological nitrogen fixation (BNF), and further promote direct N uptake (Carvalho *et al*., 2014). Within the group of soil-borne beneficial bacteria, several isolates of bacteria are collectively called Plant-Growth-Promoting Rhizobacteria (PGPR). Over the last decade a great number of PGPRs have been identified and studied in detail not only for their potential to promote plant growth, but also for their induction of biotic and abiotic stress tolerance (Vacheron *et al*., 2013; Gopalakrishnan *et al*., 2015; Verbon & Liberman, 2016; Etesami & Maheshwari, 2018). The mode of action of these PGPRs are mechanistically diverse and far from being completely understood. However, plant growth hormone-signaling, and here specifically auxin and ethylene, are likely to be involved in driving rhizobacterial effects on plant physiology. Accordingly, plant hormones modulate root development changes, such as bacteria-induced inhibition of primary root length, positive effects on lateral root formation and root hair development, as well as inhibition of cell division and cell elongation within the root apical meristem and differentiation zone (Zamioudis *et al*., 2013; Wintermans *et al*., 2016; de Zelicourt *et al*., 2018). These effects are often achieved by the secretion of plant hormone analogues directly produced by the rhizobacteria or by influencing endogenously plant growth hormone balance and signaling, causing plant developmental effects, summarized in (Tsukanova *et al*., 2017). Stimulatingly, these processes are often co-regulated by nutritional availability and nutritional signaling. For example, the PGPR *Enterobacter radicincitans* induced growth promotion and colonization increased under high-nitrogen conditions in tomato. Reciprocally, reduced N fertilization led to altered plant hormone signaling and consequently to an inhibition of these PGPR-effects (Berger *et al*., 2013). In addition, inoculation of *Arabidopsis thaliana* (Arabidopsis) with a mixture of PGPR *Bacillus spp*. induced plant growth promotion effects by enhancing N uptake and chlorophyll content, this in turn influences nutritional signaling by a bacterial-induced transcriptional induction of nitrate and ammonium transporter genes (Calvo *et al*., 2019).

Over the last decade key players regulating N-homeostasis and signaling have been identified (Wang *et al*., 2018; Fredes *et al*., 2019). These include low- and high-affinity transporter protein families such as the NITRATE-TRANSPORTER 1/PEPTIDE-TRANSPORTER FAMILY (NPF, also known as CHL1 and NRT1) and the NITRATE-TRANSPORTER 2 family (NRT2) (Krapp *et al*., 2014; O’Brien *et al*., 2016). Members of both protein families are transcriptionally regulated and functionally dependent on the plants’ nutritional N-status. For example, the high-affinity-transporting-systems *NRT2*.1 and *NRT2.2* are induced after nitrate uptake (Cerezo *et al*., 2001). Examples for root-expressed low-affinity nitrate transporters are NRT1.1 (NPF6.3) and NRT1.2 (NPF4.6) (Wang *et al*., 2018). Interestingly, it has been postulated that NRT1.1 is also able to transport auxin and consequently regulates lateral root growth in Arabidopsis (Krouk *et al*., 2010; Bouguyon *et al*., 2016). Its auxin transporting activity depends on the availability of exogenous nitrate in a concentration-dependent manner. Auxin accumulates in nitrate-rich patches of lateral roots and induces root emergence and elongation, whereas auxin depleted in sites with low nitrate availability resulting in reduced overall root growth (Remans *et al*., 2006; Mounier *et al*., 2014). Notably, root system architecture, and here primarily lateral- and primary-root length, are stimulated during moderate but homogenous nitrate availability, but inhibited under excess conditions (Jia & Wiren, 2020). This indicates that different signaling cascades are activated in fluctuating nitrate conditions resulting in adaptive physiological and morphological processes. The most studied nitrogen-signaling cascade is the so called Primary Nitrate Response (PNR), which describes the response to nitrate resupply after depletion (Medici & Krouk, 2014). This signaling pathway is rapidly activated upon nitrate perception and results in transcriptional changes within minutes. PNR is directly regulated by transcription factors like the RWP-RK transcription factor NIN-like Protein7 (NLP7) (Bellegarde *et al*., 2017). The Arabidopsis genome contains nine NIN-like Proteins (NLPs), with all of them comprising the RWP-RK motif, enabling DNA binding and transcriptional activation, and an additional PB-domain (Protein Binding) (Chardin *et al*., 2014). In addition, NLPs comprise a highly conserved nitrate-responsive domain (NRD), which determines nitrate perception and signaling transmission (Mu & Luo, 2019). In Arabidopsis, the characterization of NLP function is described for AtNLP8 and AtNLP6/AtNLP7 complex. AtNLP8 is a designated master regulator in nitrate-induced seed germination mediating catabolism of abscisic acid (Yan *et al*., 2016), while AtNLP7 functions as a master regulator in PNR (Castaings *et al*., 2009). Interestingly, NLP6 and NLP7 form heterodimers via the PB domain and are postulated to interact in complex with the transcription factor TEOSINTE

### BRANCHED1/CYCLOIDEA/PROLIFERATING CELL FACTOR1-20 (TCP20) during nitrate starvation (Guan *et al*., 2017)

Root nodule nitrogen-fixing symbiosis is evolutionary retained in different species of the NFN clade (Fabales,Rosales, Cucurbitales, Fagales), where it is established between several species and bacteria collectively named rhizobia (in Fabales and the Rosales species *Parasponia andersonii*) or Frankia, which enables them to access atmospheric dinitrogen after its fixation and conversion to ammonium by the symbiont (Werner *et al*., 2015; Griesmann *et al*., 2018). One of the central transcriptional regulators is the Nodule-INception protein (NIN), which was originally identified in *Lotus japonicus* (Schauser *et al*., 1999). Different to other NLPs, the Nitrate-responsive domain of legume NINs is only partially conserved, which could explain the loss of direct nitrate responsiveness of NIN and thus may represent an evolutional adaptation of legumes (Mu & Luo, 2019) to maintain this mutualistic interaction. However, it should be noted that the functionality of root-nodule symbiosis is strictly regulated by the availability of nitrate via NLP function as exemplified by the *Nitrate unResponsive SYMbiosis 1* (NRSYM1) in *Lotus japonicus* (Nishida *et al*., 2018). The NLP NRSYM1 accumulates in the nucleus in response to nitrate and regulates transcription of the CLE-RS2 peptide, which in turn negatively controls nodule numbers. Remarkably, the Arabidopsis AtNLP6 and AtNLP7 could partially rescue the nodulation phenotype of a *nrsym1* mutant, suggesting a functional conservation of NLPs in legumes and non-legumes plants (Nishida *et al*., 2018). In *Medicago truncatula* (Medicago), *MtNLP1* was reported to mediate nitrate signaling and to re-localize from the cytosol to the nucleus under high nitrate conditions. Subsequently, complex formation between NLP and NIN proteins in the nucleus suppresses the activation of the root-nodule symbiosis marker genes *CRE1* and *NF-YA* and thus induces the nitrate-dependent inhibition of nodulation (Lin *et al*., 2018).

Analysis of different natural soil samples indicates that the Arabidopsis rhizosphere also contains about 4% bacteria from the Rhizobiales order (Garrido-Oter *et al*., 2018) and effects of rhizobia on Arabidopsis root development have already been described. For example, the nodulating *Rhizobium sp. IRBG74* inhibits root development of Arabidopsis by impacting auxin signaling via a change in transcriptional activation of auxin responsive genes (Zhao *et al*., 2017). Similarly, *Mesorhizobium loti (M. loti)*, the natural symbiont of *L. japonicus*, inhibits Arabidopsis primary root growth in an auxin-dependent manner, whereas an auxin-independent increase in root hair formation was observed (Poitout *et al*., 2017). Furthermore, the Medicago symbiont *Sinorhizobium meliloti (S. meliloti)* induces rapid transcriptional reprogramming of defense genes in Arabidopsis cortex cells followed by late responses of stress-related genes in the pericycle, a possible stress adaptation response resulting in an increased number of lateral roots (Walker *et al*., 2017).

Here, we investigated alterations of the Arabidopsis root system architecture upon inoculation with *M. loti* and *S. meliloti* with respect to nitrate availability and nitrate signaling. We demonstrate that rhizobial effects on the Arabidopsis root architecture are initiated by nitrate signaling and specifically involve NLP-dependent pathways. Our study highlights the importance of nutrient availability during bacterial-plant interactions and shed further light on the complex interplay between plant nutritional signaling and bacterial colonization.

## Materials and Methods

### Plant growth conditions and root observations

*Arabidopsis thaliana* wild type ecotype Columbia (Col-0), Wassilewskija (WS) and knockout mutant seeds were surface sterilized with 50% ethanol/0.5% Tritron X-100 and briefly washed with 95% ethanol. Seeds for mutant plants *nlp4* (N563595), *nlp5* (N527488), and *nlp1* (N584809) were provided by *The Nottingham Arabidopsis Stock Centre* (NASC) and homozygosity for T-DNA insertion was validated by PCR with primers designed by SALK Institute Genomic Analysis Laboratory. Seeds were germinated on half-strength Murashige-Skoog-medium after stratification at 10°C for 3 days and grown horizontally for 7-days/10-days (as indicated) at 22°C with 150µmol/m^2^/s light in a 12/12-h day/night photoperiod. After germination seedlings were transferred and inoculated with bacteria on *nitrate-variable plant medium* containing 5mM/0.25mM/0.1mM (as indicated) KNO_3_ or 5mM KCl (in low-nitrate conditions KCl was used to buffer ion concentrations), 1mM KH_2_PO_4_, 0.5 mM CaSO_4_, 0.5 mM MgCl_2_, 3 mM MES, pH 5.8, 0.25% (w/v) sucrose and 0,1% micronutrients (Scheible *et al*., 2004) including 1% ferric citrate, supplemented with 0.9% Phyto-Agar and grown vertically for 3 days. Root modifications were quantified on images photographed with scale bar and analyzed on Image J Software. For root hair quantifications pictures were taken using a stereomicroscope and root hairs were measured and counted in 5 mm from the start of the differentiation zone (first root hair appearance). Data were averaged from at least 7 root samples and experiments were performed at least 3 times independently.

### Bacteria growth conditions

Rhizobium *Sinorhizobium meliloti* strain Sm2011, tetracycline resistant with mCherry as fluorescent marker, and *Mesorhizobium loti* strain MAFF, gentamycin resistant with dsRED as fluorophore, were used for bacterial treatments. For each experiment bacteria were freshly plated on TY-medium and incubated for 3 days at 28°C. Afterwards the inoculum culture was grown in 5 ml liquid TY-medium for 48h. For inoculation, bacteria were pelleted, resuspended in sterile H_2_O and diluted to a final OD_600_ of 0.1. Ten µl of bacteria solution per root were used to treat Arabidopsis seedling and primary root length was marked after inoculation. For reproducibility the timing of bacterial pre-culture growth is essential for an adequate-equal inoculum preparation.

### Quantitative RT-PCR

All quantitative RT-PCR were performed with 7-day old seedlings grown for 3 days in high- (5mM) and low- (0.25mM) nitrate conditions. Total RNA was extracted from Arabidopsis root samples using a RNA extraction kit (Sigma Aldrich) and quality control was performed photometrically using a NanoDrop (Thermo Fisher Scientific) device and on agarose gel. To remove traces of genomic DNA, 1 μg of total RNA from each sample was treated with DNase-I and genomic-DNA contamination was excluded via quantitative RT-PCR. Reverse transcription was accomplished from purified RNA with SuperScript cDNA Synthesis Kit (Quanta BioScience). Efficiency of cDNA synthesis was verified by semi-quantitative PCR on housekeeping *actin2* transcripts. Quantitative RT-PCR was performed using an *Applied Biosystems* real time PCR system and a Syber green Mix. Ct-values were normalized to the housekeeping gene *EF1α4 or UBQ4*, as indicated. Data were averaged from at least 3 independent experiments.

### FM^™^4-64 staining and image analyzes of root tip samples

Treated seedlings were incubated in 5 µM FM^™^4-64 dissolved in liquid *nitrate-variable plant medium* for 15 min. After 5 min washing in liquid plant medium primary root tips were mounted on a microscopy slide. Images of 3880×3880 resolution were taken with SP8-Leica confocal microscope using a 25x water objective and a white laser, emission of 554 to 591 nm.

### Phylogenetic analyzes

NLP sequences were retrieved by the tBLASTn algorithm v2.9.0+ (Camacho *et al*., 2009) using the NIN protein from the model species *Medicago truncatula* as query, a database containing 124 land plants genomes from the SymDB database (http://www.polebio.lrsv.ups-tlse.fr/symdb/web/) and no e-value greater than 1^e-10^. Sequences were aligned using MAFFT v7.407 (Katoh & Standley, 2013) with default parameters. Alignments were cleaned using trimAl v1.4 rev22 (Capella-Gutierrez *et al*., 2009; Guindon *et al*., 2010) to remove positions with more than 50% of gaps. Cleaned alignments were subjected to Maximum Likelihood analysis using IQ-TREE v1.6.7 (Nguyen *et al*., 2015) after testing the best-fitting evolutionary model using ModelFinder (Kalyaanamoorthy *et al*., 2017). Branch support was tested with 10,000 SH-aLRT (Guindon *et al*., 2010). Trees were visualized and annotated using the iTOL platform v4.4.2 (Letunic & Bork, 2019). For readability reasons, presented phylogeny in (Fig. S2) has been reduced to 7 species with maintained topology of the 124 species-main tree (Fig. S1).

### Accession numbers

Sequence data from this article can be found under the following Arabidopsis Genome Initiative accession numbers: NRT2.1 (AT1G08090), NRT1.1 (AT1G12110), NIA1 (AT1G77760), NIR1 (AT2G15620), RSL4 (AT1G27740), EXT11 (AT5G49080), RHD6 (AT1G66470), CYCD3;1 (AT4G34160), CYCB1;1 (AT4G37490), UBQ4 (AT5G25760), EF1α4 (AT5G60390), NLP1 (AT2G17150), NLP4 (AT1G20640), NLP5 (AT1G76350), NLP6 (AT1G64530), NLP7 (AT4G24020)

## Results

### Rhizobia associate with the root surface of Arabidopsis seedlings

Following the findings that rhizobia and other diazotrophic bacteria are able to colonize the endosphere of non-legume roots by entering via natural openings such as sites of lateral root emergence (de Zelicourt *et al*., 2018; Schneijderberg *et al*., 2018) we assessed the colonization patterns of rhizobia on Arabidopsis roots. For this, Arabidopsis plants were grown vertically *in vitro* and inoculated with fluorescently labelled *M. loti* for three days. Bacterial biofilms formed along the root epidermis (Fig. 1a) with enrichments being observed at the root cap and the root elongation zone (Fig. 1b). Furthermore, rhizobia accumulated at sites of lateral root emergence (Fig. 1c) and invaded the space of these natural epidermal openings (Fig. 1d). However, clear signs of intracellular, endophytic colonization were not observed under these conditions. From this point of view rhizobia rather associated with the non-legume Arabidopsis by establishing an interaction on the outside of the plant root, than infect as endophytes plant tissue.

**Figure 1.:**
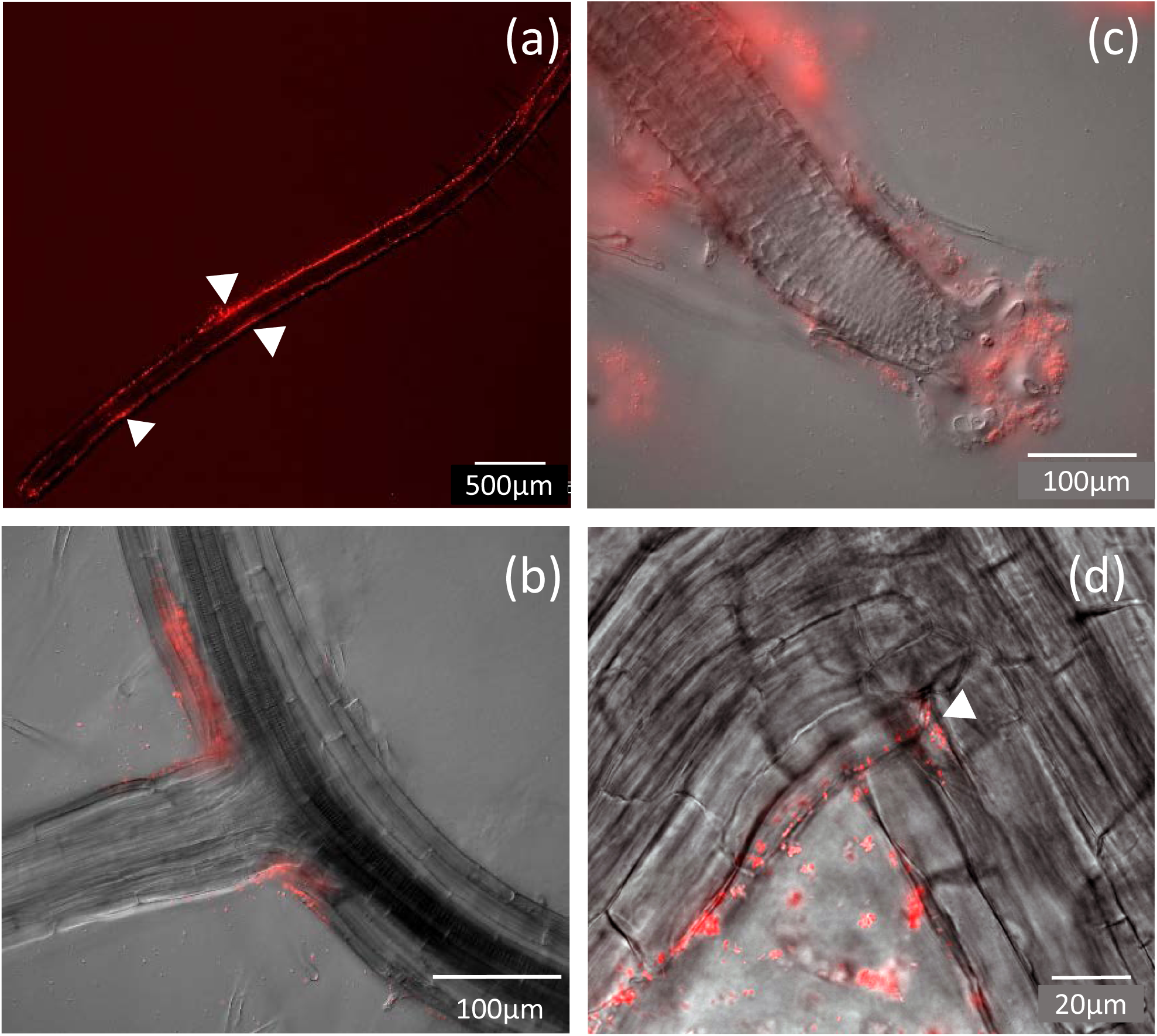
Rhizobia attachment on roots of non-legume host plant Arabidopsis. 10-days old Arabidopsis seedlings inoculated with transgenic rhizobia (*M. loti*) tagged with dsRed fluorophore grown on plant media with 5mM nitrate. (a) Bacterial biofilm formation on growing Arabidopsis primary root (white arrows indicate bacterial enrichments (hot spots)). (b) Bacteria accumulation on root tip and root elongation/differentiation zone. (c) and (d) *M. loti* accumulation at lateral root emergence and intercellular space invasion, validated by z-stack. Samples were observed under Zeiss stereo fluorescence microscope with mCherry filter Ex/Em: 587/647 nm as well as for (d) confocal-microscope Leica SP8.

### Rhizobia affect plant root development under high nitrate conditions

As several soil-borne bacteria are known to alter the plant root architecture, thus enabling plants to better access nutrients directly or indirectly, we tested whether *M. loti* and *S. meliloti -*effects depend on the availability of exogenous nitrate. For this, we transferred 10-days old Arabidopsis seedlings onto plant medium with high nitrate (5mM) and nitrate starvation (0.1mM KNO_3_) conditions and additionally inoculated these plants with either *S. meliloti* or *M. loti* for 3 days. The number and length of lateral roots significantly increased when inoculating plants grown under high-nitrate conditions with rhizobia compared to the non-inoculated control (Fig. 2a-e) while the primary root length was unaffected by this treatment (Fig. 2f). Nitrate deprivation fully abolished the rhizobial effect on lateral root growth (Fig. 2d, e) and only resulted in a moderate increase in the primary root length, which was independent of the presence of bacteria (Fig. 2f). Thus, in our conditions we could not observe reported negative effect of rhizobia on primary root length in Arabidopsis (Zhao *et al*., 2017).

**Figure 2.:**
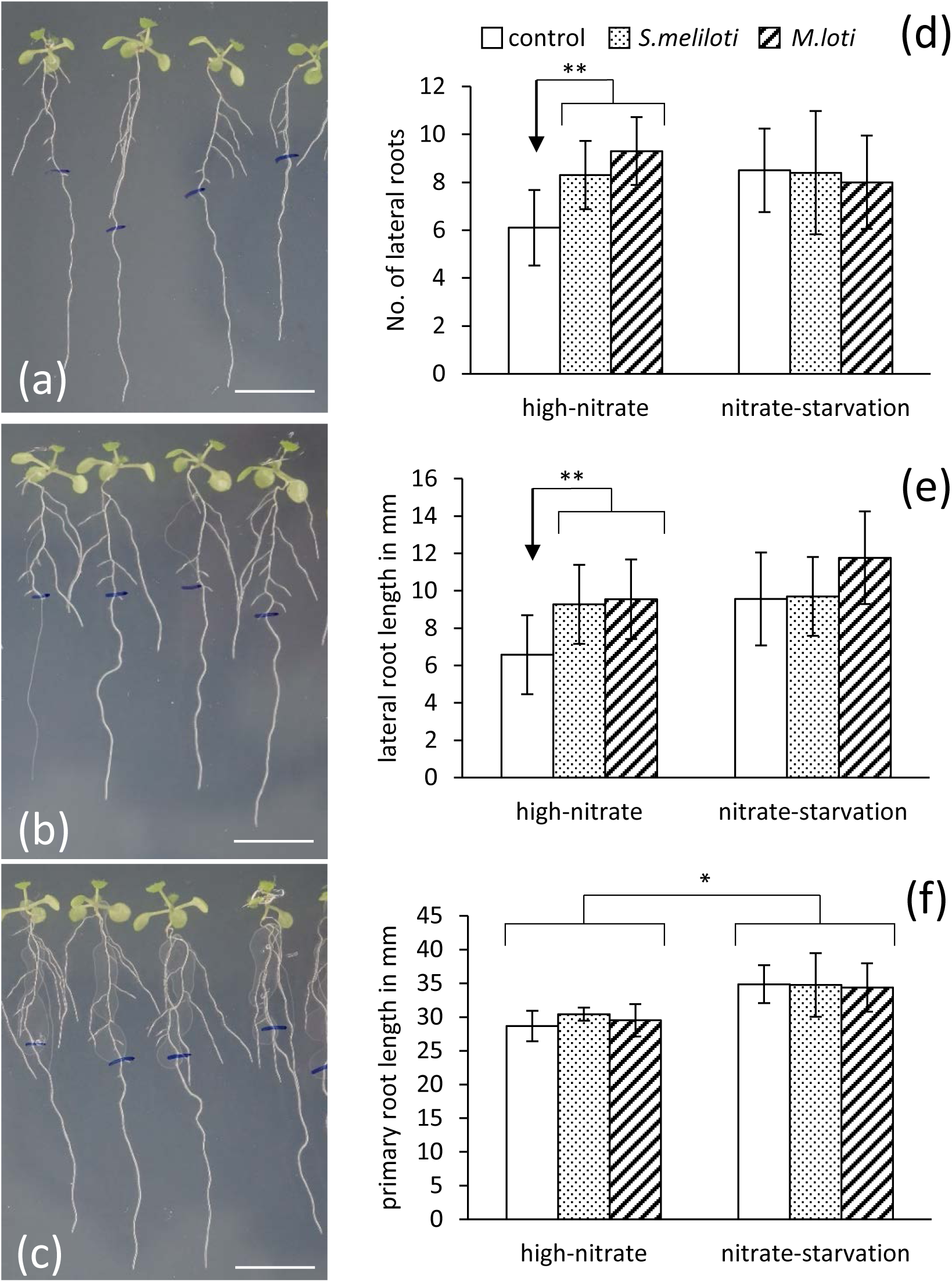
Rhizobia-induced modifications on Arabidopsis root architecture depend on nitrate availability. Ten-days old seedlings were transferred on high-nitrate plant medium (5mM KNO_3_) or nitrate-starvation plant medium (0.1mM KNO3) and roots were inoculated for 3 days with rhizobia (*S. meliloti*) or (*M. loti*). (a) control seedlings grown for 3 days on high-nitrate plant medium. (b) *S. meliloti* and (c) *M. loti* inoculated seedlings grown on high-nitrate plant medium scale bar presents 1 cm (starting root length, before inoculation, was marked with blue marker pen). (d)-(f) Quantifications of No. of lateral roots, lateral root length, and primary root length on different nitrate conditions and indicated rhizobia treatment. Statistical significance was determined by students t-test *=p<0.05; **=p<0.01.

A primary contact point of rhizobacteria and plants are root hairs. As another described plant modifying effect of rhizobia is to alter root hair formation (Poitout *et al*., 2017; de Zelicourt *et al*., 2018), we investigated root hair development under these two nitrate conditions. As expected, nitrate starvation led to a significant increase in root hair length (Fig. 3a-b, g) and number (Fig. 3h)(Vatter *et al*., 2015) even in the absence of rhizobia. While the presence of *S. meliloti* only moderately affected root hair patterns (Fig. 3c-d, g-h) under control conditions, a significant induction of root hair growth was observed upon inoculation with *M. loti* under elevated nitrate conditions. This bacterial effect was largely lost under nitrogen-starvation conditions (Fig. 3e-h). To further test the impact of rhizobia on root hair growth, we examined root hair-specific marker gene expression in roots treated with rhizobia under different nitrate availability conditions. Indeed, rhizobia tended to induce the morphogenetic root-hair genes ROOT HAIR DEFECTIVE6 (*RHD6*), RHD6-LIKE4 (*RSL4*), and EXTENSIN11 (*EXT11*) (Fig. 3i-k), thus *M. loti* treatments were particularly efficient. Overall, these data demonstrate that Arabidopsis root architecture and morphology are affected by rhizobia in a nitrate-dependent manner.

**Figure 3.:**
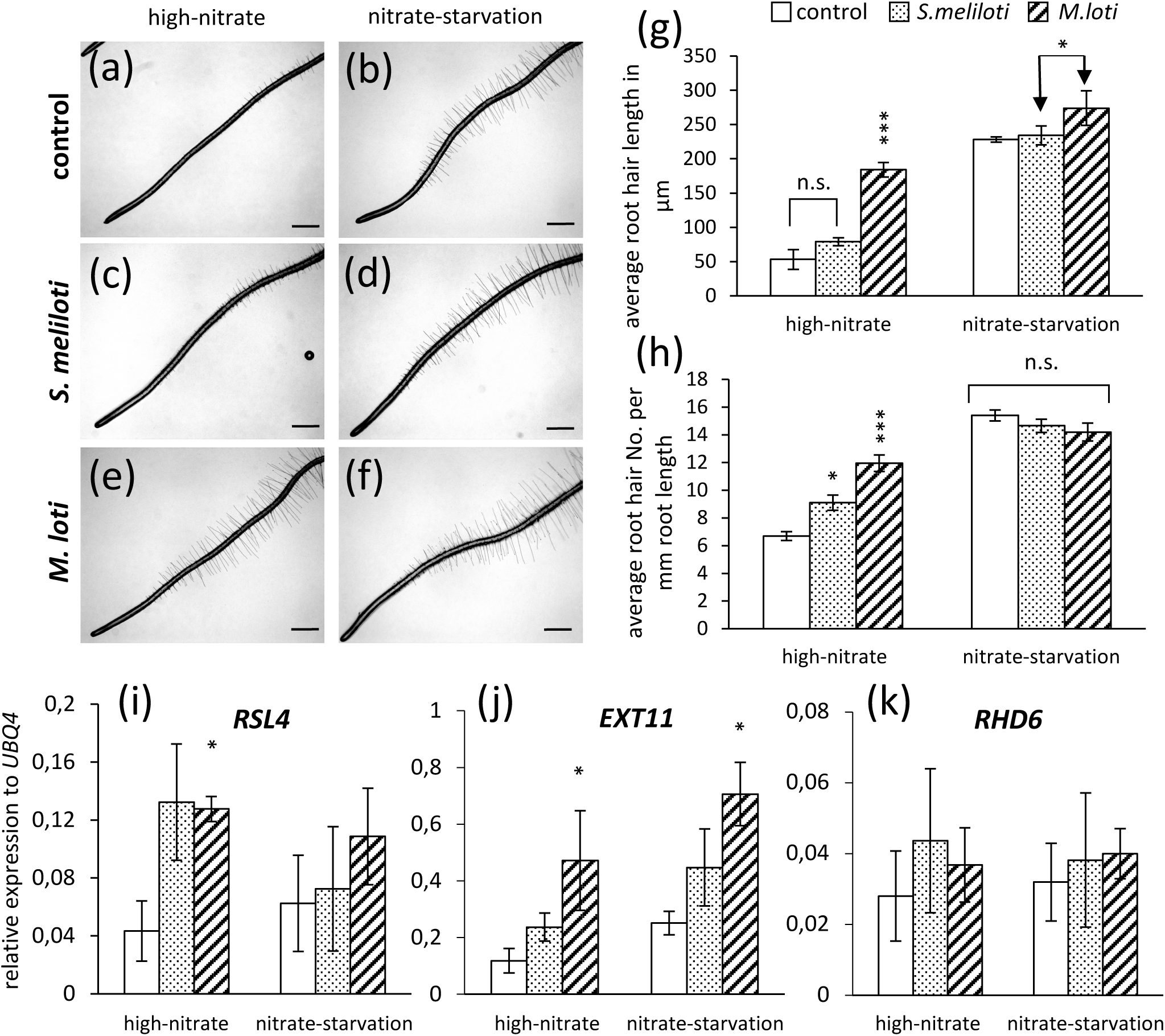
Rhizobia-induced root hair formation depends on high-nitrate availability. Seven-days old seedlings were transferred on high-nitrate plant medium or nitrate-starvation plant medium and roots were inoculated for 3 days with rhizobia (*S. meliloti*) or (*M. loti*). Primary root tip of control samples after 3 days on plant medium (a) on high nitrate condition and (b) starvation condition, (c), (d) *S. meliloti* treated samples, and (e), (f) *M. loti* treated samples, scale bar presents 100µm. (g) and (h) quantification of root hair formation on primary root tip (average of first 5mm). (i)-(k) Transcript levels of indicated root hair-related genes, specifically ROOT HAIR DEFECTIVE6 (*RHD6*), RHD6-LIKE4 (*RSL4*), and EXTENSIN11 (*EXT11*), quantified by qRT-PCR and calculated relative to the housekeeping gene expression level of *UBQ4*. Statistical significance was determined by students t-test *=p<0.05; ***=p<0.001 and n.s. = not significant

### Rhizobia influence nitrate-related gene expression

Next, we tested whether rhizobia-induced root modifications depend on nitrate signaling responses. Therefore, we investigated nitrate-related gene expression in roots treated with rhizobia. Since transcriptional reprogramming differ in low-nitrate conditions (limiting condition) compared to nutrient depletion conditions (harsh condition), and as we wanted to focus on nitrate-induced signaling rather than depletion stress, we decided to change our low-nitrate conditions to 0.25mM KNO_3_. The selection of specific marker genes was based on the regulation mechanisms of NLP7 (primary-nitrated-response) (Marchive *et al*., 2013). Among them are transceptors *NRT*s that have been reported to be up-regulated in low-nitrate conditions and to improve nitrate use efficiency. Although *NRT2.1* and *NRT1.1* were induced in low-nitrate conditions, their expression was further increased in samples treated with rhizobia, for *NRT1.1* even in high-nitrate conditions (Fig 4a,b). On the other side, genes involved in nitrate metabolism such as *NIA1* (nitrate reductase), and *NIR1* (nitrite reductase) which are regulated during nitrate response, were found to be induced upon rhizobia inoculation of plants grown under high-nitrate, but not under low nitrate conditions (Fig. 4c-d). This suggests that rhizobia differentially regulate nitrate-related genes in Arabidopsis and influence nitrate signaling mainly in high-nitrate conditions.

**Figure 4.:**
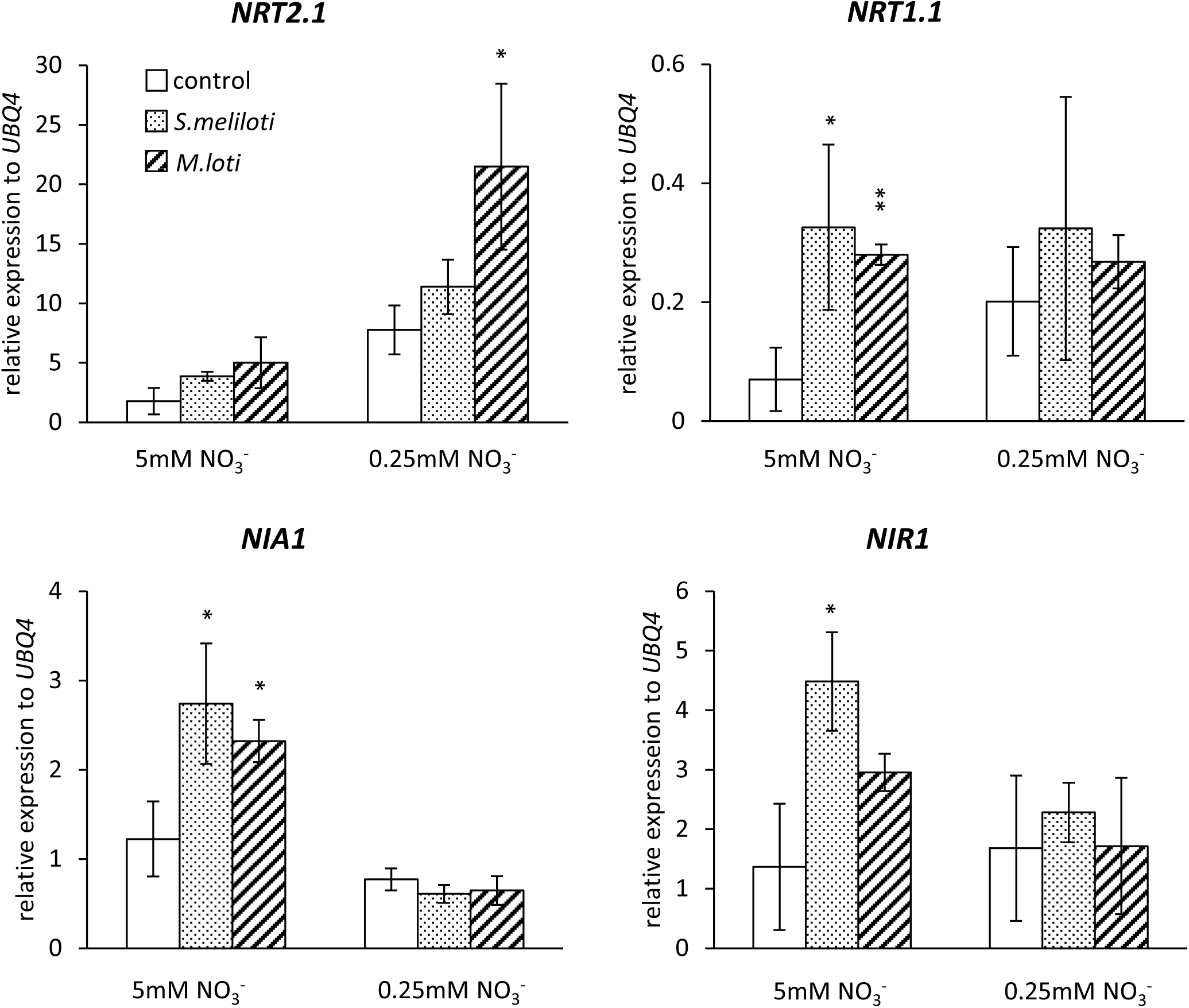
Rhizobia-induced expression of nitrate-regulated genes. Transcript levels of indicated genes in 7-day old Arabidopsis root samples treated for 3-days with rhizobia *S. meliloti, M. loti* or in 10-day old control samples grown on high-(5mM) or low-(0.25mM) nitrate plant medium conditions. Quantified by qRT-PCR and calculated relative to the housekeeping gene expression level of *UBQ4*. Statistical significance was determined by students t-test *=p<0.05; **=p<0.01

### *M. loti*-induced root morphology changes are controlled by NLP4 and NLP5

As expression patters of nitrate-related genes in *M. loti*-treated root samples could indicate a nitrate-dependent signaling during *M. loti*-induced root morphology changes, we further tested the genetic dependency of our observations on root development using different nitrate signaling mutants. Recently, it was shown that NIN-like proteins play a role in nitrate signaling during rhizobia-legume interactions. These proteins suppress gene expression by either directly binding of NIN-proteins or competitive binding promotors of symbiotic-responsive genes (Lin *et al*., 2018; Nishida *et al*., 2018). These roles of NLPs during nitrate-dependent signaling prompted us to genetically assess their importance during Rhizobia-Arabidopsis interactions. Large-scale phylogenetic analysis of the NLP family revealed their conservation in land plants (Schauser *et al*., 2005). Furthermore, different duplication events led to three NLP-clades that evolved prior to the divergence of seed plants (Fig. S1 and S2).

As NLP7 (NIN –like protein 7) was identified as key regulator during the primary nitrate response in Arabidopsis and functions in complex with NLP6 and TCP20 in nitrate signaling (Guan, 2017), we obtained independent t-DNA insertion mutant alleles for NLP6 and NLP7 (Castaings *et al*., 2009). Under control conditions (uninoculated, high-nitrate condition) *nlp6* and *nlp7* mutants already developed more root hairs compared to the wild-type, while mutants responded to the presence of rhizobia similar to the wild-type in a nitrate-independent manner (Fig. 5a-b). As we could not explain rhizobia-induced nitrate signaling on genetic control of the NLP7/NLP6 complex by the observed mutant phenotypes, we decided to investigate the NLP-clade NLP1-NLP5 (closely related to the clade containing NIN from legumes such as Medicago) (Fig. S2). Like WT plants, t-DNA insertion mutants of this NIN-related gene family in Arabidopsis, specifically *nlp1, nlp4* and *nlp5*, responded to nitrate limitation by increased root hair formation (Fig. S3, Fig. 5c-d). However, and in contrast to the *nlp1* mutant, *nlp4* and *nlp5* mutants failed to induce pronounced root hair development upon application of *M. loti* (Fig. 5c-d, Fig. S4a-b). Furthermore, the rhizobia effects of induced lateral root emergence and overall lateral root length were also lost in *nlp4* and *nlp5* mutants (Fig. S4, Fig. 5c-d). These data suggest the presence of an NLP-dependent signaling pathway in Rhizobia-Arabidopsis interactions, similar to nitrate-dependent regulation of symbioses.

**Figure 5.:**
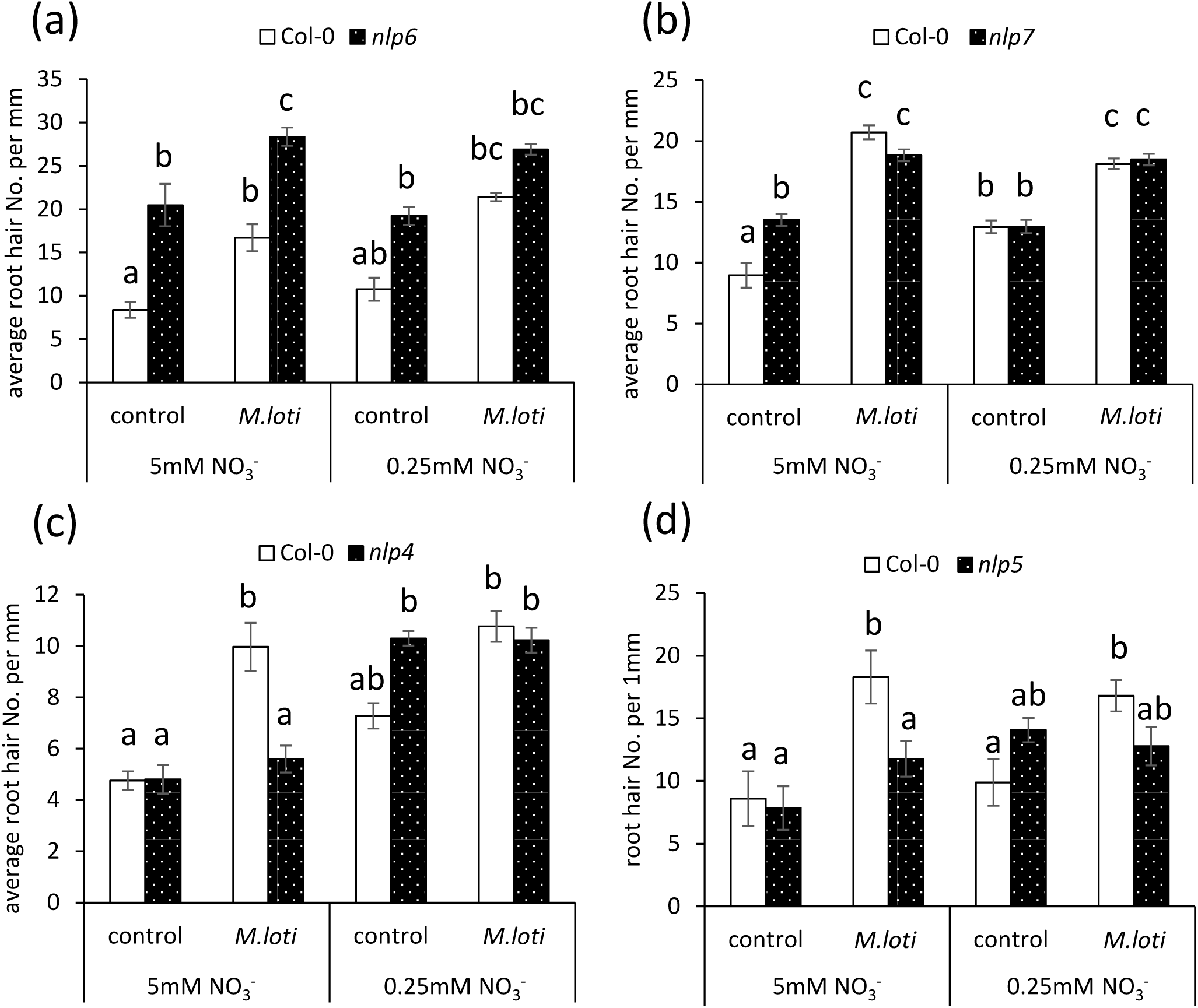
Specific NIN-like Proteins are involved in *M. loti* induced root hair effect. Seven-days old seedlings of wild-type (Col-0) or mutant plants (as indicated) were transferred on high-nitrate plant medium (5mM KNO_3_) or low-nitrate plant medium (0.25 mM KNO_3_) and roots were inoculated for 3 days with *M. loti*. Quantification of root hair formation on primary root tip is presented. Statistical significance was determined by one-factorial ANOVA α=0.05 followed by Scheffé post-hoc test

### *M. loti*-triggered effects on root hair patterning depend on the transceptor NRT1.1

Nitrate is sensed and transported via specific membrane proteins comprising high- and low-affinity transceptors, which are functionally dependent on nitrate availability. The best studied transporter protein is the dual-affinity transceptor NRT1.1, which changes its affinity to nitrate depending on its phosphorylation status (Sun & Zheng, 2015). On the other side, an example for an interesting high-affinity transporter is NRT2.1, which is also involved in the development of lateral roots and is transcriptionally regulated upon several environmental stimuli including external nitrate availability (Orsel *et al*., 2002; Little *et al*., 2005). Due to expression profiles of NRT-genes in roots treated with rhizobia (Fig. 4a,b), we studied *M. loti* induced root modifications in *nrt1.1-5* (*chl1-5*) and *nrt2.1* mutant backgrounds. In analogy to *nlp4* and *nlp5* mutants, the *nrt1.1* mutant also entirely failed to induce root hair formation upon inoculation with *M. loti* under high- and low-nitrate conditions (Fig. 6a). Application of *M. loti* did not affect root hair length in the *nrt1.1* mutant (Fig. 6c). In contrast to the *nrt1.1* mutant, the *nrt2.1* mutant fully responded to the presence of rhizobia, with even more and longer root hairs forming on this genotype (Fig. 6b,d). These results indicate an important role for NRT1.1 in rhizobia-induced root modifications.

**Figure 6.:**
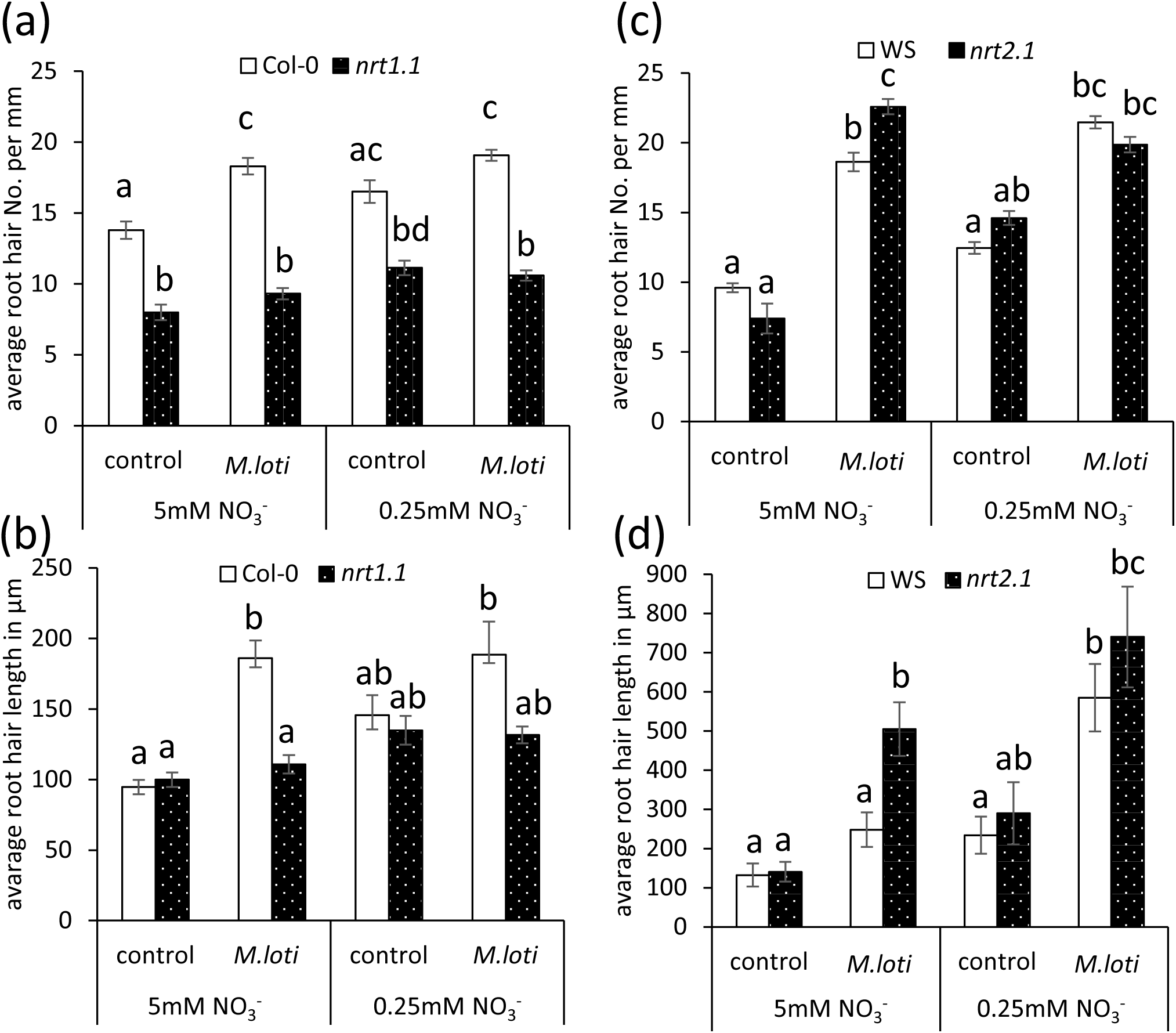
Dual-affinity-transceptor *NRT1.1* is needed for *M. loti*-induced impact on root morphology, but not *NRT2.1*. Seven-days old seedlings of wild-type (Col-0) and Wassilewskija (WS) or mutant plants (as indicated) were transferred on high-nitrate plant medium (5mM KNO_3_) or low-nitrate plant medium (0.25mM KNO_3_) and roots were inoculated for 3 days with *M. loti*. Quantification of root hair formation and root hair length on primary root tip (first 5mm) is presented. Statistical significance was determined by one-factorial ANOVA α=0.05 followed by Scheffé post-hoc test.

### *M. loti* alters cell proliferation and elongation patterns in the root tip

Besides the effects of nitrate as a signaling component on developmental aspects of root development it also influences cell divisions at the root apical meristem and subsequent cell cycle progression (Guan *et al*., 2017; Gigli-Bisceglia *et al*., 2018). Similar effects were also described for treatments with PGPRs, which can influence root architecture via hormonal cross-talks (Verbon & Liberman, 2016). Since the perception of rhizobial signals by legume roots results in the onset of an organogenesis program, we tested possible impacts of *M. loti* on Arabidopsis root cell patterning. For this, we transferred 7 days-old Arabidopsis seedlings on plant medium with high- or low-nitrate, inoculated them with *M. loti* for 3 days and stained root cell membranes with the dye FM^™^4-64 to visualize cell morphology and quantify cell numbers (Fig. 7a-d). In wild-type Arabidopsis plants, the average length of the first five cortex cells above the root transition zone significantly decreased under high-but not under low-nitrate conditions in the presence of *M. loti* (Fig. 7e), whereas this effect was not observed in the *nlp4* and *nlp5* mutant plants (Fig. 7f,g). This shorted elongation zone could suggest a bacteria-induced early differentiation, dependent on NLP4 and NLP5, in high-nitrate conditions, and could explain the observed effects on root hair density on the primary root tip (Fig. 3h and 5c,d). For the quantification of meristem size, we counted longitudinally the cell number from the quiescent center until the transition zone (first doubling of cell size) in the cortex cell layer. Cortical cell numbers were found to be induced by *M. loti* in both wild-type and *nlp4* mutant plants, but this rhizobial effect was lost only in *nlp5* mutants (Fig. 7h-j). This is accompanied by increased transcription of the cell cycle-related cyclin D3 (*CYCD3;1*) (Fig. 7h), but not of *CYCB1;1* (Fig. 7i). Since *CYCD3;1* is expressed during the meiotic G1/S phase of the cell cycle, while *CYCB1;1* serves as a marker for a prolonged G2 phase, we conclude that cortical cell division rates are induced by *M. loti*. Overall, we demonstrate that rhizobia-induced signaling influences cell cycle progression and root cell elongation in a NLP5-dependent manner.

**Figure 7.:**
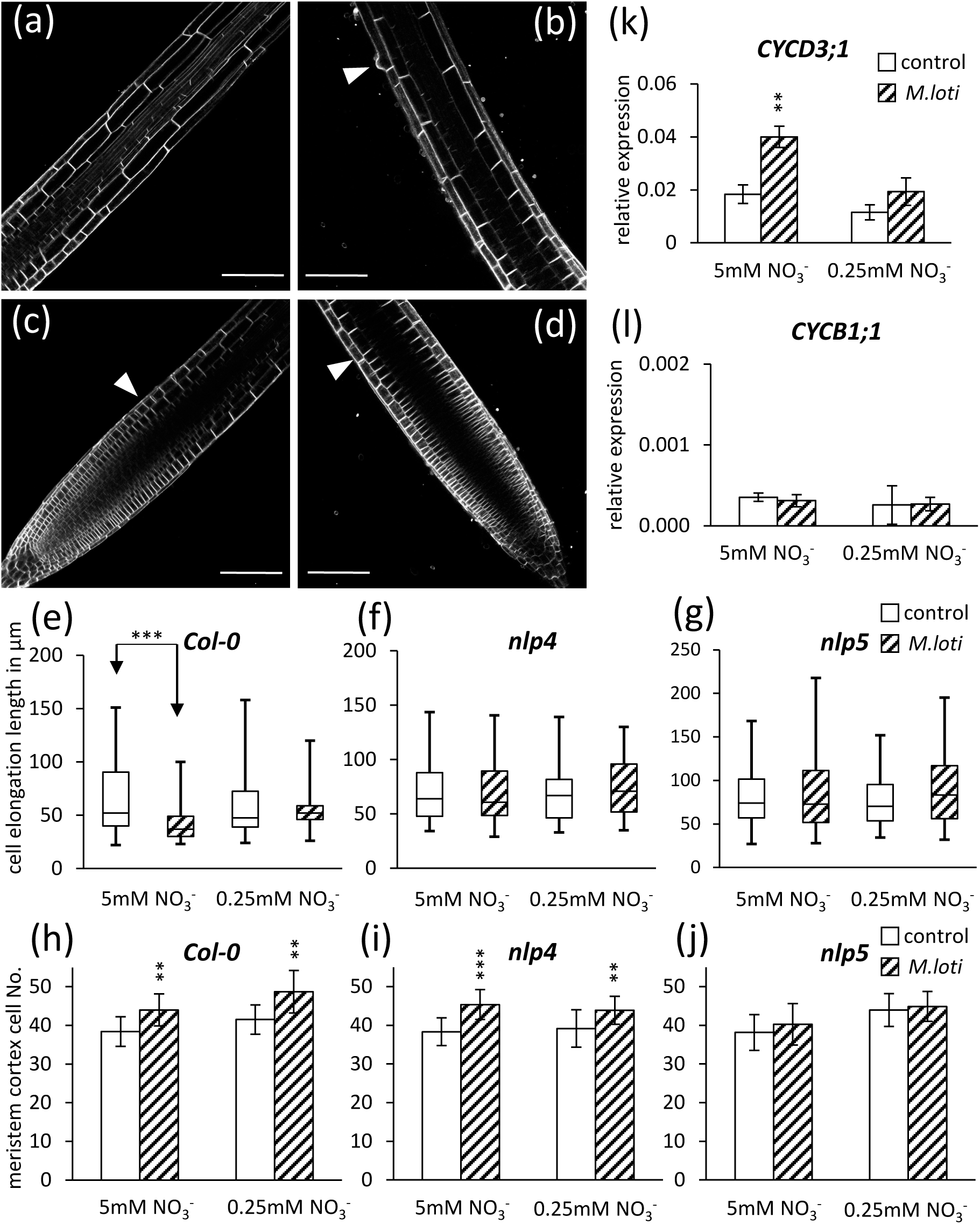
*M. loti* impacts cell- proliferation and –elongation. Seven-days old seedlings of wild-type (Col-0) or indicated mutant plants were transferred on high-nitrate plant medium or low-nitrate plant medium and roots were inoculated for 3 days with *M. loti* (control = no inoculation). (a)-(j) FM_™_ 4-64 staining of primary root tip and visualization at SP8 Leica confocal microscope. (a) and (b) representative pictures of first cells of root elongation zone (white arrow indicates first root hair cell), (c) and (d) pictures of primary root apical meristem (white arrow indicates end of meristem), (a, c) control and (b, d) *M. loti* on high-nitrate samples, white arrows show cell size change, scale bar presents 100 µm. (e) - (g) length of the first 5 elongated cells, of primary root samples with or without *M.loti* grown on high- or low- nitrate plant medium. (h) – (j) quantification of meristem cortex cell no. on primary root tip. (k) and (l) transcript levels of indicated genes quantified by qRT-PCR and calculead relative to the housekeeping gene expression level of *EF1α4*. Statistical significance was determined by students t-test **=p<0.01; ***=p<0.001

## Discussion

Gene regulatory networks control plant responses to different nutrient conditions, leading for instance to modification of the root architecture. In natural soils, plant roots are additionally challenged by a great diversity of beneficial and pathogenic microbes and the plants’ overall nutrient status determines the responsiveness to these microbes (Shahzad & Amtmann, 2017; Singh *et al*., 2019). Among them are rhizobia, which have been mostly investigated in frame of the root-nodule symbiosis, a process that ultimately results in the ability of these bacteria to fix atmospheric dinitrogen gas. This mutualistic interaction strictly depends on the unavailability of nitrate and is immediately terminated upon its resupply of inorganic nitrate to the host plant (van Noorden *et al*., 2016). While the root-nodule symbioses interaction mostly relies on a highly strain-specific interaction with a narrow host range, a microbiome study revealed the presence of rhizobia in the rhizosphere or as endophytes in roots of non-legume plants including Arabidopsis (Garrido-Oter *et al*., 2018).

In this manuscript, we investigated rhizobia-induced root morphology changes and their dependence on nitrate availability and nitrate-related signaling cascades in Arabidopsis. We demonstrate that rhizobia affect lateral root emergence and -elongation (Fig. 2) and increase root hair density and length (Fig. 3, 5). Interestingly, these effects were found to be nitrate-dependent and abolished under nitrate starvation conditions (Fig. 2, 3) implying an underlying signaling crosstalk between nitrate-related signaling and rhizobia-induced responses.

It is well documented that nitrate starvation is accompanied by stimulated lateral root emergence and primary root elongation, while excess nitrate leads to root growth inhibition (Jia & Wiren, 2020). In line with this, the presence of PGPRs on Arabidopsis roots stimulates a low-nitrate response. This can be exemplified by the PGPR *Phyllobacterium* STM196 that positively influences root architecture under high-nitrate conditions. These rhizobacteria did not directly alter the N-status in leaves or root nitrate uptake, but induced the expression of nitrate starvation marker genes such as *AtNRT2.5* and *AtNRT2.6* (Mantelin *et al*., 2006) that are also postulated to play a central role in bacterial-induced growth promotion (Kechid *et al*., 2013). This is in agreement with our results of *M. loti*-induced root hair development that were more pronounced under high-nitrate conditions (Fig. 3, 5). While it cannot be excluded that these and other bacteria partially metabolize nitrogen sources in the rhizosphere, thus subsequently reduce nutrient availability for the plant, our data indicate a rather specific nitrate-related signaling circuit that steers rhizobial-effects on the Arabidopsis root system (Fig. 4). This predominantly involves the transcription factors *NLP4* and *NLP5*, as the corresponding mutants failed to morphologically respond to the presence of rhizobia (Fig. 5, 7, Supp. 3). This is of particular interest, considering the evolutionary proximity of AtNLP4 and AtNLP5 to the legume gene NIN, and their co-orthologs legume genes *MtNLP1* and *LjNLP4* that are essential for controlling nitrate-dependent root nodule symbiosis (Schauser *et al*., 2005; Lin *et al*., 2018; Nishida *et al*., 2018; Mu & Luo, 2019). Thus, the response to rhizobia could be trait shared within the NLP clade (NLP1-5, angiosperms) and the Eudicots duplication leading to NLP4/5 and NLP1/2/3, which finally allowed the possible neofunctionalization of NIN for root nodule symbiosis (Fig. S2). Furthermore, we demonstrated an NLP4/NLP5-dependent and rhizobium-induced stalling of cell elongation but increased cell division rates at the proximal part of the root elongation zone (Fig. 7), which suggests a NLP-dependent (nitrate related) -signaling in the root elongation zone, direct to and earlier differentiation of roots treated with rhizobia. In line with this hypothesis, cell wall integrity and meristem cell progression are postulated to depend on nitrate reductase function and endogenous cytokinin homeostasis in roots tips (Gigli-Bisceglia *et al*., 2018). In addition, morphological changes also coincided with an enhanced activation of the cell-cycle marker gene *CYCB1;1* (Trinh *et al*., 2018). Reciprocally, the application of *Pseudomonas putida WCS417* on Arabidopsis resulted in an inhibition of root growth, which was likely caused by reduced cell elongation and driven by an observed activation of *CYCB1;1* in inoculated samples (Zamioudis *et al*., 2013). While *M. loti* induced similar effects on cell proliferation we observed an induced expression of *CYCD1;3* but not of *CYCB1;1* indicative for an accelerated cell progression into the G1 phase (Fig. 7). Such induction of D-like cyclins (here: CYCD1, D2 and D3) was also reported in rice seedlings when treated with *S. meliloti* (Wu *et al*., 2018).

The importance of nitrate-regulated transcription factors during plant root development has been extensively described. For example, the bZIP transcription factors TGA1 and TGA4 control nitrate-induced lateral root formation (Alvarez *et al*., 2014) and root hair cell fate, latter by regulating the root hair cell-specific gene *CAPRICE* (*CPC)* (Canales *et al*., 2017). Furthermore, a meta-analysis indicates nitrate regulation of root hair growth by bZIP and G2-transcription factors coordinated auxin signaling response (Canales *et al*., 2014). Interestingly, the NRT1.1-TGA1/4 nitrate signaling complex additionally controls root hair density and trichoblast cell length (Canales *et al*., 2017). This is in agreement with our data, showing that NRT1.1 also controls rhizobia-induced differences in trichoblast patterning (Fig.6). We additionally observed a differential up-regulation of the nitrate transporters *NRT1.1* and *NRT2.1* under low-nitrate conditions, which was further pronounced in the presence of rhizobia (Fig. 4). A similar effect was observed for *NRT2.1* upon inoculation of Arabidopsis and *Lactuca sativa* roots with the PGPR *Pseudomonas nitroreducens* (Trinh *et al*., 2018).

Taken together we understand the investigated bacterial-plant interaction as a complex signaling pathway between nutritional sensing followed by metabolic progressing in crosstalk with bacteria interaction and stress-induced responses, which finally results in developmental adaptation processes. Conscious of the well documented crosstalk between nitrate signaling and growth hormones (Takei *et al*., 2004; Vidal *et al*., 2010; Ma *et al*., 2014; Naulin *et al*., 2020), we consider observed rhizobia-induced nitrate responses as early signaling events after bacterial perception, which finally leads to downstream fine-tuning of growth hormone effects (Fig. 8). Nitrate-controlled signaling in plant-microbe interaction is at the beginning of its investigations, though direct new perspectives to understand and decipher environmental adaptation processes of plants. In the future, these findings will facilitate the development of bacterial inoculums as effective and sustainable solutions in agriculture.

**Figure 8.:**
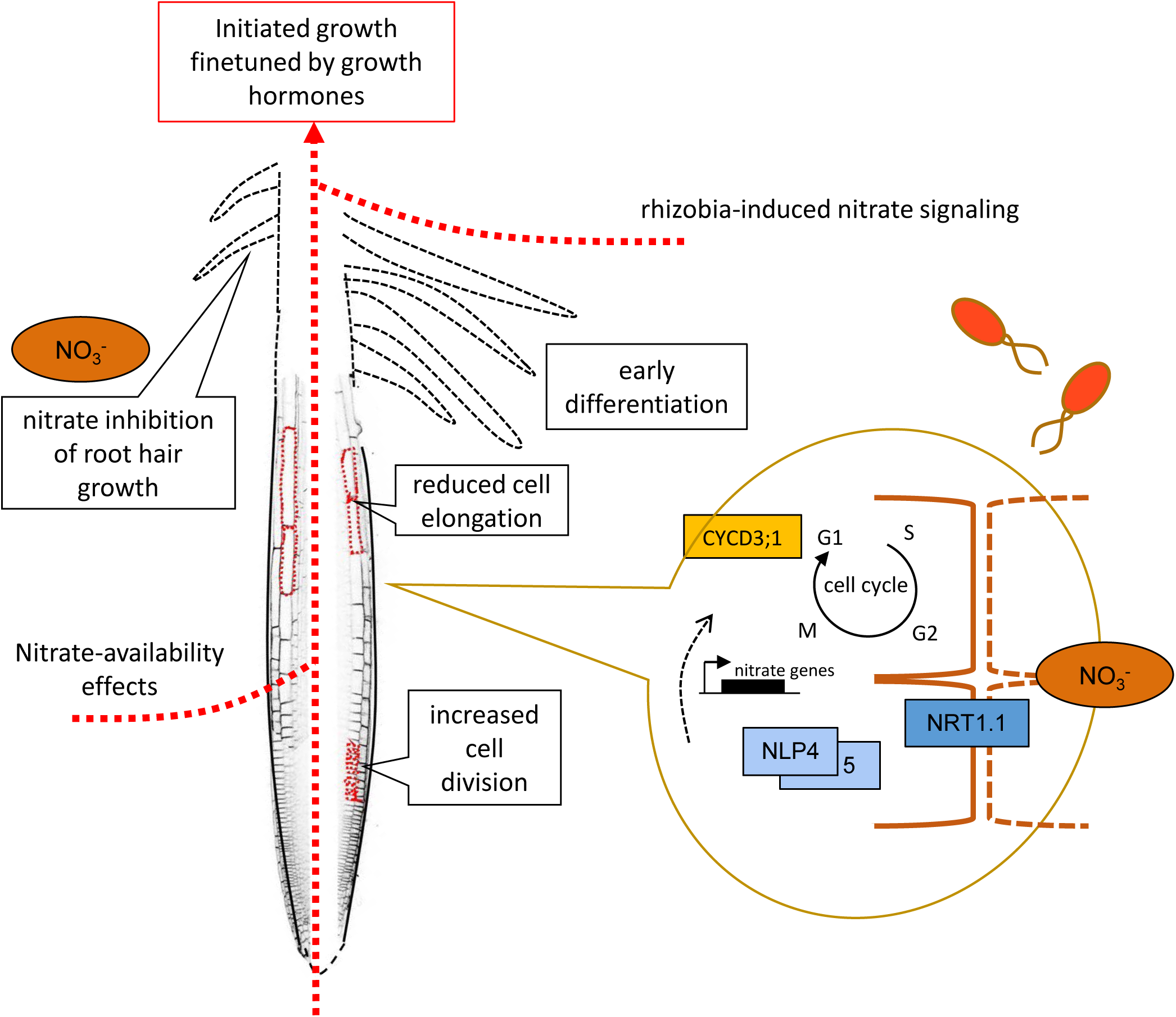
Summary of rhizobia-induced nitrate signaling in roots of *Arabidopsis thaliana*. Results obtained indicate rhizobia-induced local root development effects on the non-legume host *Arabidopsis thaliana*. We demonstrate an involvement of nitrate-related key proteins and show an induction of nitrate signaling during rhizobia responses, which finally impacts in cell cycle progression and morphological modifications. Furthermore, described effects are dependent on nitrate availability and suggest a complex cross-talk between bacterial- and nutritional- induced signaling pathways.

## Acknowledgements

Authors thank Liên BACH for mutant plants *chl1-5* (*nrt1.1*) and Anne Krapp for kindly providing seeds for *nrt2.1-1, nlp6.2* and *nlp7*.1. This study was supported by the German Research Foundation (Deutsche Forschungsgemeinschaft) under Germany’s Excellence Strategy (CIBSS– EXC-2189 – Project ID 39093984). This work was partly supported through a grant to the University of Cambridge by the Bill & Melinda Gates Foundation (OPP1172165) and UK government’s Department for International Development (DFID). The Laboratoire de Recherche en Sciences Végétales (LRSV) laboratory belongs to the TULIP Laboratoire d’Excellence (ANR-10-LABX-41). Especially, we thank all members of the *OttLab* team for the active discussions and supporting ideas that helped to develop the project.

## Author Contribution

STS and TO conceived and designed the experiments; STS, EL, JK performed the experiments; STS, JK, P-MD, analyzed and interpreted the data; and STS, JK, P-MD and TO wrote the manuscript.

## Supporting Information

Fig. S1 Phylogenetic tree of NIN-like Proteins (main tree).

Fig. S2 Reduced phylogenetic tree of NIN-like Proteins.

Fig. S3 NLP1 mutation is not influencing Rhizobia effects on root hair development.

Fig. S4 AtNLP4 and AtNLP5 are involved in *M. loti* induced root modifications.

Table S1 Primer sequences used in this study

## Notes

### Competing Interest Statement

The authors have declared no competing interest.

