## Supporting Information for "Integration of A Nitrate-Related Signaling Pathway in Rhizobia-Induced Responses During Interactions with Non-Legume Host Arabidopsis thaliana"

**Figure S1.: Phylogenetic tree of NIN-like Proteins (main tree).** Maximum Likelihood tree of the NLP family (model: GTR+F+R10; log-likelihood: -598245.4704). Tree is rooted on the non-vascular plants. Clades are colored and named according the *Arabidopsis thaliana* nomenclature. Only significant branch support values (>80%) are shown.

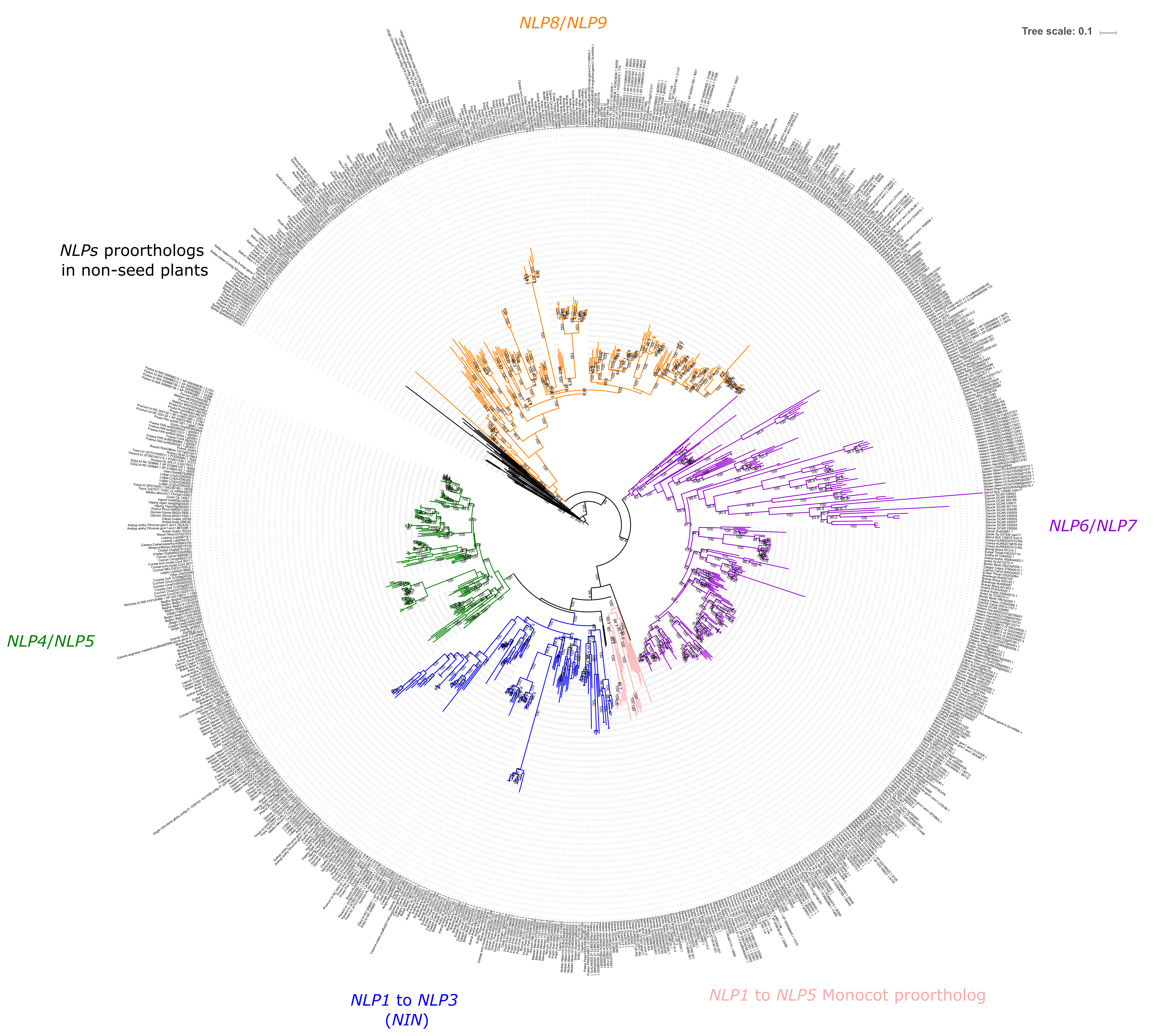
NLP8/NLP9
Tree scale: 0.1
NLPs proorthologs
in non-seed plants
NLP6/NLP7
NLP4/NLP5
NLP1 to NLP3
(NIN)
NLP1 to NLP5 Monocot proortholog

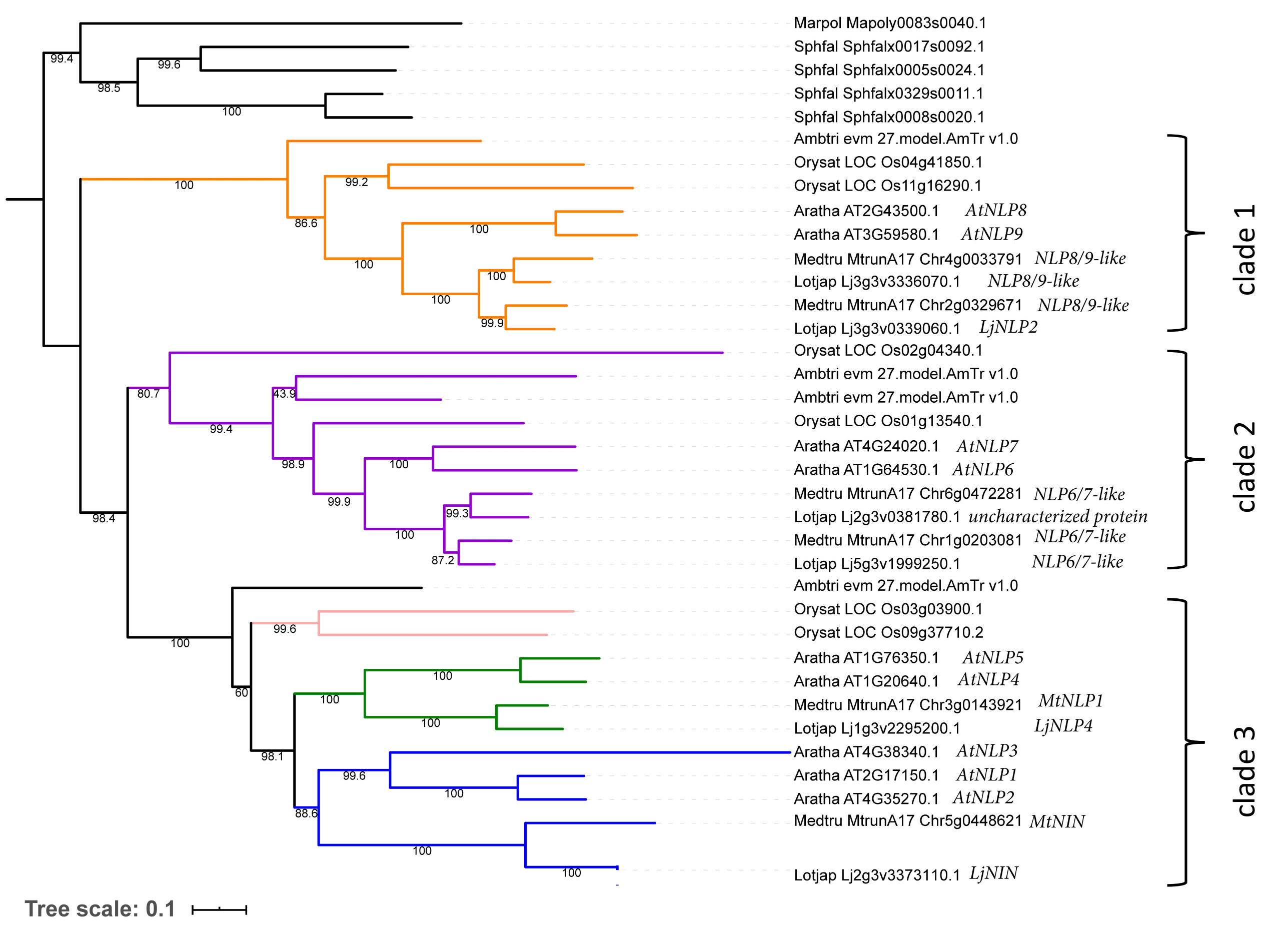


**Figure S2.: Reduced phylogenetic tree of NIN-like Proteins.** NLP sequences were retrieved using the tBALSTn algorithm v2.9.0+ by the NIN protein from the model species *Medicago truncatula* as query. Reduced likelihood tree of the NLP family. Clades are colored and named according the *Arabidopsis thaliana* nomenclature. Only significant branch support values (>80%) are shown.

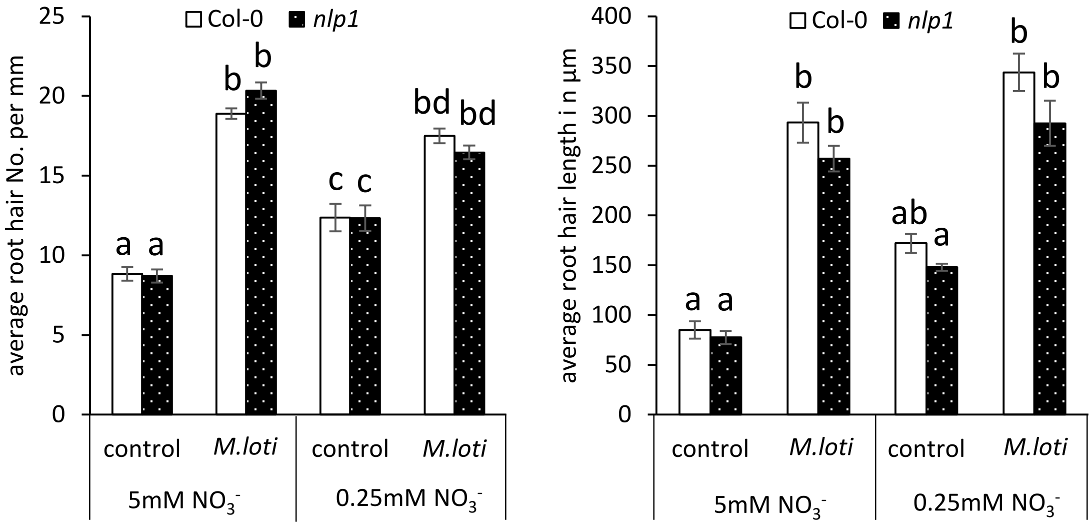


**Figure S3.: NLP1 mutation is not influencing rhizobia effects on root hair development.** Seven-days old seedlings of wild-type (Col-0) or mutant plants (as indicated) were transferred on high-nitrate plant medium (5mM $KNO_3$) or low-nitrate plant medium (0.25mM $KNO_3$) and roots were inoculated for 3 days with M. loti. Quantification of root hair formation and root hair length on primary root tip (first 5mm) is presented. Statistic significance was determined by one-factorial ANOVA α=0.05 followed by Scheffé post-hoc test

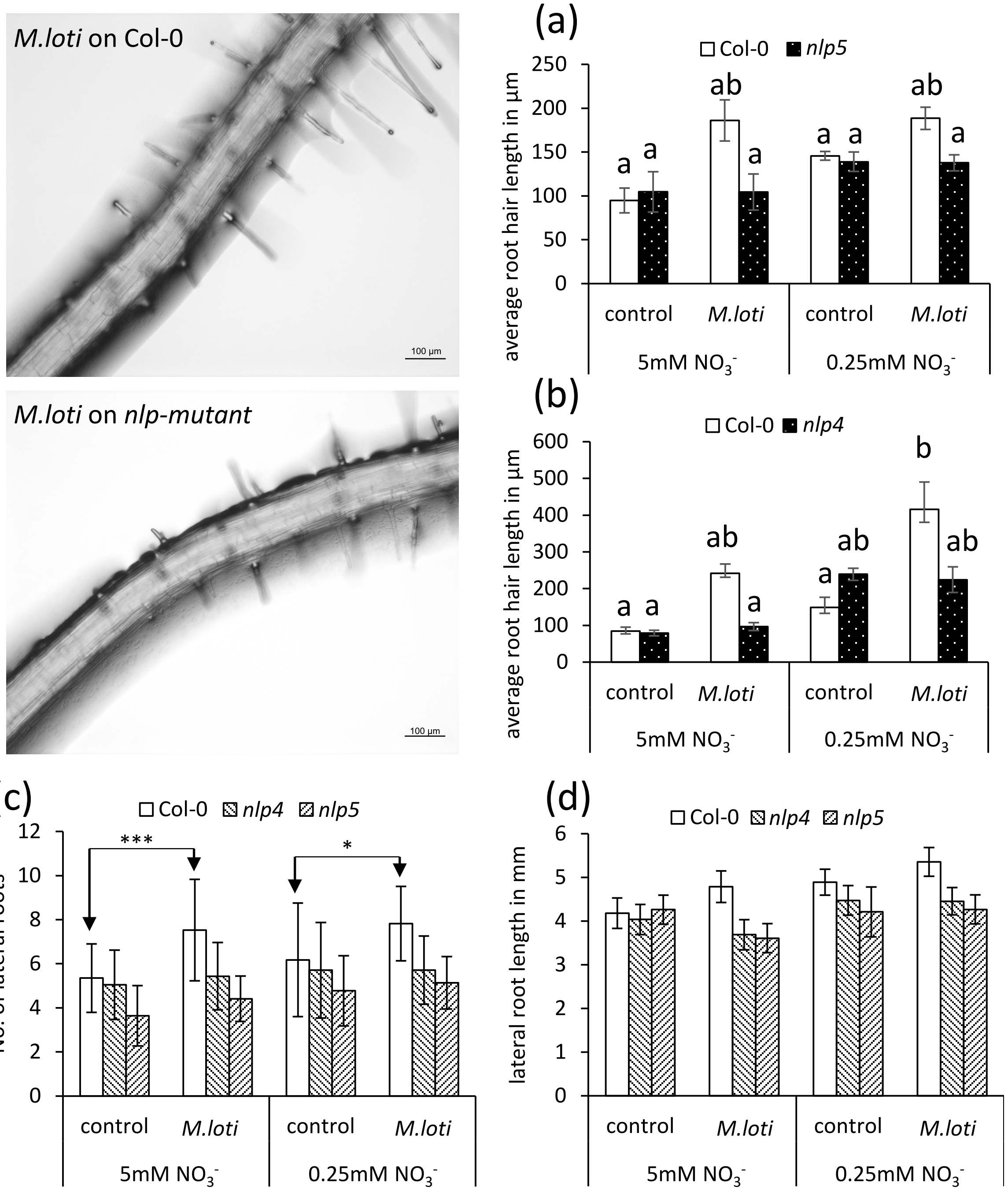


**Figure S4.: AtNLP4 and AtNLP5 are involved in *M. loti* induced root modifications.** Seven-days old seedlings of wild-type (Col-0) or mutant plants (as indicated) were transferred on high-nitrate plant medium (5mM $KNO_3$) or low-nitrate plant medium (0.25 mM $KNO_3$) and roots were inoculated for 3 days with *M. loti.* (a) and (b) quantification of root hair length on primary root tip (first 5mm) is presented. Left site representative close-up image of Col-0 and nlp-mutant (*nlp5*) root differentiation zone treated with M. loti for 3 days. Statistic significance was determined by one-factorial ANOVA α=0.05 followed by Scheffé post-hoc test. (c) and (d) quantifications of lateral root No. and lateral root length is presented. Statistic significance was determined by students t-test *=p<0.05; ***=p<0.001.

**Table S1 Primer sequences used in this study**

| | |
|---|---|
| RSL4_F | AACCTTGTGCCAAACGGGAC |
| RSL4_R | CCAGGCCGTTGTAAGCCAAT |
| EXT11_F | GGCAGTTTTTGGTTTATGTCGTG |
| EXT11_R | GATGGCTGGGAAAATTGTATTGTG |
| RHD6_F | GCCCTAGATCCACCGAAACTCC |
| RHD6_R | TGGCTGCTAGGCTTTGTGG |
| NRT1.1_F | GCACATTGGCATTAGGCTTT |
| NRT1.1_R | CTCAATCCCCACCTCAGCTA |
| NRT2.1_F | AGTCGCTTGCACGTTACCTG |
| NRT2.1_R | ACCCTCTGACTTGGCGTTCTC |
| NIA1_R | ACGGAGCATGGATGAGTT |
| NIA1_F | ATCGTCAAAGAAACCGAAGTC |
| NIR1_F | TGCTGATGACGTTCTTCCACTCTGC |
| NIR1_R | CTGAGGGTTGACTCCGAAATAGTCTC |
| CYCB1;1_F | TCAGCTCATGGACTGTGCAA |
| CYCB1;1_R | GATCAAAGCCACAGCGAAGC |
| CYCD3;1_F | CTTCAGCTCGTTTCTGTCGC |
| CYCD3;1_R | TGTCTCCTCCACTTGAAAGTCT |
